# Calcium-based input timing learning

**DOI:** 10.1101/2024.11.14.623617

**Authors:** Shirin Shafiee, Sebastian Schmitt, Christian Tetzlaff

## Abstract

Stimulus-triggered synaptic long-term plasticity is the foundation of learning and other cognitive abilities of the brain. In general, long-term synaptic plasticity is subdivided into two different forms: homosynaptic plasticity describes synaptic changes at stimulated synapses, while heterosynaptic plasticity summarizes synaptic changes at non-stimulated synapses. For homosynaptic plasticity, the Ca^2+^-hypothesis pinpoints the calcium concentration within a stimulated dendritic spine as key mediator or controller of underlying biochemical and -physical processes. On the other hand, for heterosynaptic plasticity, although theoretical studies attribute important functional roles to it, such as synaptic competition and cooperation, experimental results remain ambiguous regarding its manifestation and biological basis. By integrating insights from Ca^2+^-dependent homosynaptic plasticity with experimental data of dendritic Ca^2+^-dynamics, we developed a mathematical model that describes the complex temporal and spatial dynamics of calcium in dendrites.We show that the influx of calcium into a stimulated spine can lead to its diffusion to neighboring spines, triggering heterosynaptic effects such as synaptic competition or cooperation. By considering different input strengths, our model explains the ambiguity of reported experimental results of heterosynaptic plasticity, suggesting that the Ca^2+^-hypothesis of homosynaptic plasticity can be extended to also model heterosynaptic plasticity. Furthermore, our model predicts thata synapse can modulate the expression of homosynaptic plasticity at a neighboring synapse in an input-timing-dependent manner, without the need of postsynaptic spiking. The resulting sensitivity of synaptic plasticity on input-spike-timing can be influenced by the distance between involved spines as well as the local diffusion properties of the connecting dendritic shaft, providing a new way of dendritic computation.

## 1 Introduction

Long-term synaptic plasticity is a key mechanism for learning and memory in the brain (Caya-Bissonnette and Beique 2024; Lynch 2004) and inspiration for modern artificial neural network algorithms (Bellec et al. 2020; Kasai et al. 2021), with the majority of studies focusing on homosynaptic plasticity, describing synaptic changes of stimulated synapses or dendritic spines. On the other hand, experimental studies have revealed that the induction of synaptic plasticity exerts its influence not only on the spine undergoing stimulation, but also extends to affect neighboring spines (Chistiakova et al. 2015; Hayama et al. 2013; Jenks et al. 2021). As shown by theoretical studies, such heterosynaptic influences are required for network stabilization or synaptic competition (Bannon et al. 2020; Chistiakova et al. 2014; Chistiakova and Volgushev 2009; Jenks et al. 2021). Different to homosynaptic plasticity, the biological underpinnings of heterosynaptic plasticity and its detailed implementation are still to a large degree unknown (Chater and Goda 2021).

Focusing on homosynaptic plasticity, the majority of theoretical and experimental investigations have highlighted the significance of the intracellular calcium concentration in a dendritic spine as a critical determinant of synaptic strength modulation, summarized in the Ca^2+^-hypothesis; a high level of [Ca^2+^] induces long-term potentiation (LTP), a medium level of [Ca^2+^] triggers long-term depression (LTD), while a low level does not result to significant synaptic changes (Graupner and Brunel 2012; Hiratani and Fukai 2017; Lisman 1989; Luboeinski and Tetzlaff 2021; Shouval et al. 2002; Wang et al. 2005a). Hereby, calcium functions mainly as a secondary messenger, initiating and modulating signaling cascades that involve for instance phosphatases and kinases triggering synaptic plasticity (Graupner and Brunel 2012).

In recent years, studies show that dendrites hosting spines have a more active role than a passive cable that sums synaptic signals and transmits them to the soma (Poirazi and Papoutsi 2020; Stuart et al. 2016). For instance, dendrites integrate electrical signals in a non-linear manner (Poirazi and Mel 2001; Yuste 2023) or can detect sequences of inputs (Bhalla 2017; Branco et al. 2010) such that they are considered as biochemical units, playing also an important role in synaptic plasticity (Nishiyama and Yasuda 2015; Yasuda 2017). Furthermore, *in-vitro* and *in-vivo* experiments show that correlated synaptic inputs cause an elevation in calcium concentration in the dendrite (Gidon et al. 2020; Harvey and Svoboda 2007; Inglebert and Debanne 2021; Lee et al. 2016; Nevian and Sakmann 2006; Wang et al. 2005b) and strong inputs can even lead to the release of Ca^2+^ from local dendritic stores (Emptage et al. 1999; Mahajan and Nadkarni 2019; Martucci and Cancela 2022; Nishiyama et al. 2000; Padamsey et al. 2019; Royer and Paré 2003).

In this study, we have developed a first computational model integrating the well-established Ca^2+^-hypothesis of synaptic plasticity and recent results of dendritic Ca^2+^ dynamics (Fig. 1A). We demonstrate that, upon stimulation, the elevated Ca^2+^ concentration within one spine can lead to the diffusion of Ca^2+^ ions through the dendritic shaft to neighboring spines. Depending on the state of the neighboring spine, the diffused calcium might cause dramatic alterations in the pattern of homo- and heterosynaptic plasticity, resulting to a new type of learning rule that yields synaptic changes depending on the difference in timing of spikes of neighboring spines, without the need of postsynaptic action potentials. To understand the resulting complex Ca^2+^ and plasticity dynamics, we systematically varied stimulation protocol, number of spines, and Ca^2+^-related parameters of spines and the dendrite. First, we show in a system consisting of two spines being connected by a dendrite, the emergence of heterosynaptic competition and cooperation dependent on the frequency of input spikes. Subsequently, we considered larger systems consisting of a multitude of spines situated on a dendritic shaft and unified the results from different experimental data sets (Chater et al. 2022; Oh et al. 2013; 2015; Tong et al. 2021) by considering different levels of input frequency. Hereby, we could verify our assumption of Ca^2+^ diffusion as key regulator for heterosynaptic plasticity. Considering the two-spine system, finally we show that calcium-based homo- and heterosynaptic plasticity could result to spike-timing-sensitive synaptic changes. Similar to the well-established spike-timing-dependent plasticity protocols for homosynaptic plasticity (Graupner and Brunel 2010; 2012), we observe various complex types of “temporal windows” of potentiation and depression, providing the basis of a new type of learning rules highlighting the potential role of heterosynaptic plasticity for computation and cognition.

**Figure 1.**
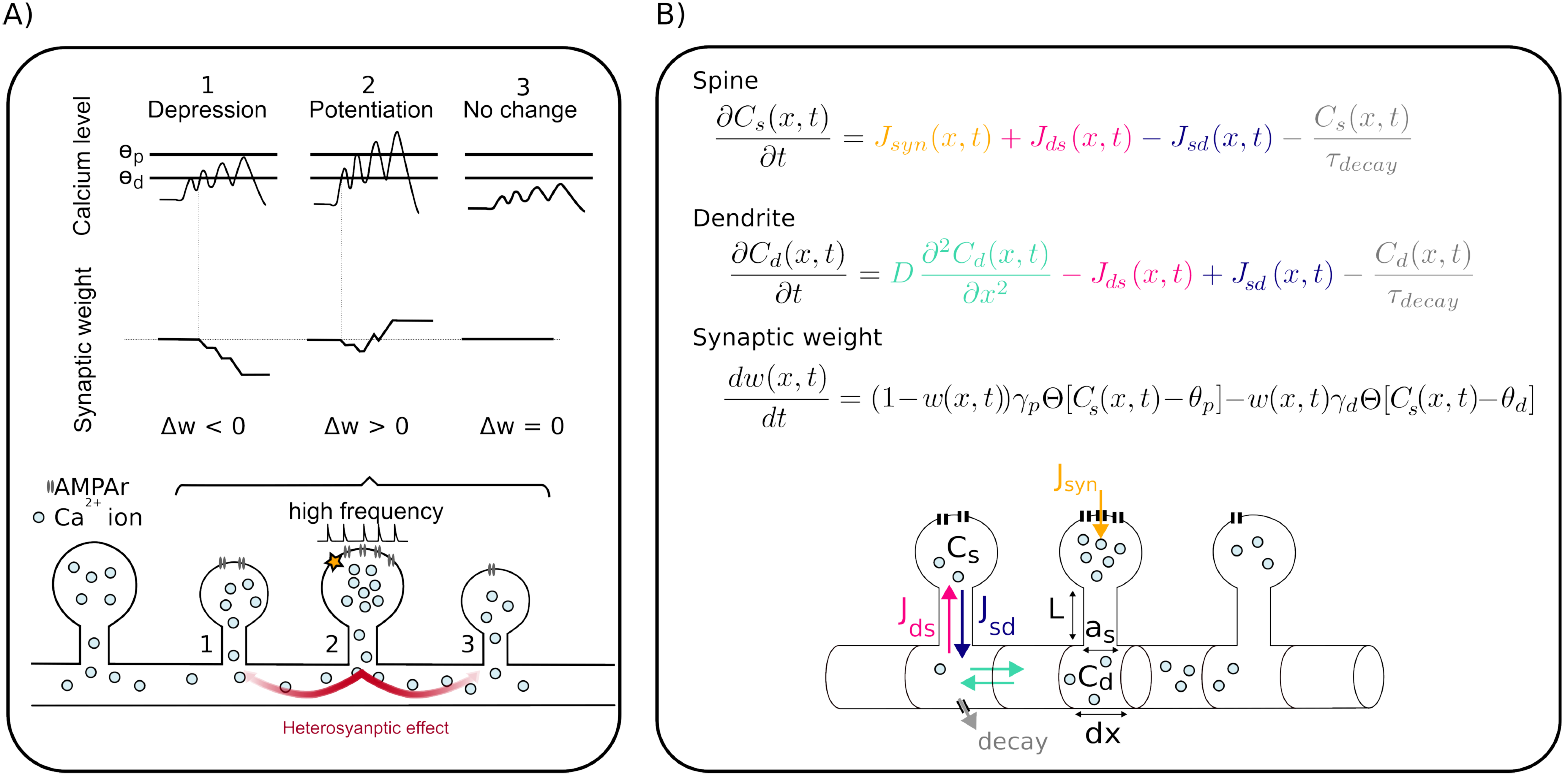
Illustration of our model of calcium diffusion between spines. **A**) Schematic of main mechanisms in our model given the example of three neighboring spines: The stimulated spine (#2) experiences potentiation, while the calcium level of spine #1 could only rise slightly and undergoes depression. A modest amount of calcium in spine #3 does not substantially alter the synaptic weight. **B**) Mathematical model of the calcium dynamics and resulting synaptic weight changes. For more details see Methods section.

## 2 Model and Methods

We introduce a biophysical computational model of the dynamics of the concentration of free Ca^2+^ ions in the dendritic shaft and dendritic spines, and resulting synaptic weight changes (Fig. 1B). We consider the dendrite as a tube of radius *r*_*d*_ that we divide into many small segments, each of length *dx*. For different setups, a different number of “average” spines is being situated at some dendritic segments. If we assume a spine at segment *x*, the diffusion equation governing the change of the concentration of free calcium within the dendritic shaft is modeled by the following equation:

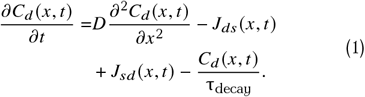

*C*_*d*_ (*x, t*) represents the calcium concentration (in µM) at the specified time *t* and segment *x*. The first term on the right hand side describes the spatial diffusion of the dendritic calcium concentration. The last term on the right hand side summarizes the loss of freely diffusing calcium by binding to proteins or flux into the extracellular space with time constant τ^decay^. If segment *x* does not host a spine, the terms *J*_*ds*_ and *J*_*sd*_ are set equals zero. Otherwise, calcium flux from the dendritic segment to the spine is defined by *J*_*ds*_ (*x, t*), and *J*_*sd*_ (*x, t*) is the flux from spine to the dendritic compartment, following:

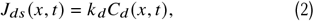

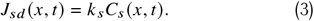

Rates *k*_*d*_ and *k*_*s*_ are given by the inverse of the mean first passage time for influx (see Biess et al. 2007; Biess et al. 2011) to the spine and to the dendrite respectively:

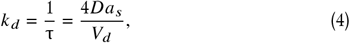

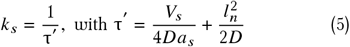

with 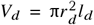 and 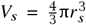 being the volume of the dendritic segment and spine, respectively. *r*_*s*_ and *r*_*d*_ are the radius of spine and dendritic compartments, *l*_*d*_ is the length of dendritic compartment, and *l*_*n*_ is the length of spine neck (see Table 1).

**Table 1:** Used parameters.

| Parameter | Symbol | Value | Source |
| --- | --- | --- | --- |
| Diffusion coefficient of free $\text{Ca}^{2+}$ | $D$ | $220 \frac{\mu\text{m}^2}{\text{s}}$ | Allbritton et al. 1992; Means et al. 2006; Biess et al. 2011 |
| Time constant of calcium leakage | $\tau_{\text{decay}}$ | 0.08 s | Yasuda 2017 |
| Neck length | $l_n$ | $0.5 \mu\text{m}$ | Araya 2014 |
| Neck radius | $a_s$ | $0.1 \mu\text{m}$ | Araya 2014 |
| Spine radius | $r_s$ | $0.34 \mu\text{m}$ | Araya 2014 |
| Dendrite radius | $r_d$ | $1 \mu\text{m}$ | O'Donnell et al. 2011 |

Calcium concentration in the spine at dendritic segment *x* is as follows with the same leak constant τ_decay_ as the dendrite:

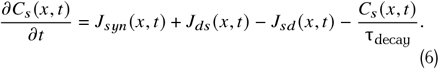

In spines, the synaptic electrical current, resulting from a presynaptic spike, yields a calcium influx according to

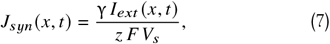

where *I*_*ext*_ (*t*) is the synaptic input current, *F* is Faraday constant and *z* is the calcium valance. γ is the fraction of electrical current converted to calcium current, which is reported to be 0.11 (Biess et al. 2011).

As presynaptic input, we consider a spike train with a mono-exponential decay:

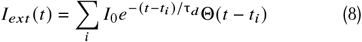

with *I*_0_ = 0.1 pA, τ_*d*_ = 1 ms, and presynaptic spike times *t*_*i*_. Following the calcium hypothesis of (homo)synaptic plasticity (Graupner and Brunel 2012; Shouval et al. 2002), we consider that a high calcium concentration *C*_*s*_ in the spine, which can be caused by high input rates, induces potentiation and strengthens the synaptic weight, while medium level of calcium induces depression and weakens synaptic strength. Following Graupner and Brunel 2012, we use a threshold-based plasticity model where the weight of the synapse or spine at segment *x* is updated based on the level of its calcium concentration *C*_*s*_:

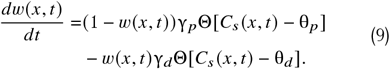

Here θ_*p*_ and θ_*d*_ are thresholds for potentiation and depression, Θ is the Heaviside function, γ_*p*_ and γ_*d*_ are LTP and LTD constants. When calcium concentration is above θ_*p*_ or θ_*d*_, potentiation or depression occurs respectively (Fig. 1A).

Numerical simulations were conducted by discretizing the diffusion partial differential equation temporally and spatially, implementing an implicit (backward) Euler scheme for solution convergence and stability (see Supplementary methods for more details).

## 3 Results

Using our computational model of the dynamics of freely diffusing Ca^2+^ ions (Fig. 1), first, we considered a piece of dendrite, at which two similar spines are situated with a distance of 1µ*m*. We inject a synaptic current at one of these two spines (spine 1, blue in Fig. 2A), while the other spine remains unstimulated (spine 2, orange). Synaptic activity results in a high calcium concentration and an accumulation of calcium ions at the stimulated spine (Fig. 2A, middle), leading to potentiation (right). Subsequently, free calcium ions diffuse to adjacent dendritic compartments and, by this, to the non-stimulated spine. Here, diffusion yields a mildly increased calcium level in spine 2 such that the non-stimulated synapse undergoes a slight depression (orange, Fig. 2A, right). This pattern of homo- (spine 1) and heterosynaptic (spine 2) plasticity is called synaptic competition (Chater et al. 2022; Chistiakova et al. 2015; Jenks et al. 2021). Fig. 2B shows the influence of high frequency stimulation on the magnitude and direction of plasticity. In contrast to a single spike stimulation, subjecting spine 1 to a burst of stimulations causes a more pronounced elevation of calcium level at the neighboring spine located in proximity to the stimulated spine (orange, Fig. 2B). This elevated calcium concentration results in potentiation of both the stimulated spine and the adjacent unstimulated spine, introducing heterosynaptic cooperation (Chen et al. 2013; Chistiakova et al. 2015). Fig. 2C illustrates another protocol similar to the first one in Fig. 2A. We initially stimulated the first spine (blue), but different to the previous protocol, here, this stimulation is followed by a stimulation of the second spine (orange) 15ms later. Compared to the first protocol, stimulation of the second spine completely changes the pattern of plasticity, triggering synaptic depression at spine 1 and potentiation of spine 2 (Fig. 2C, right).

**Figure 2.**
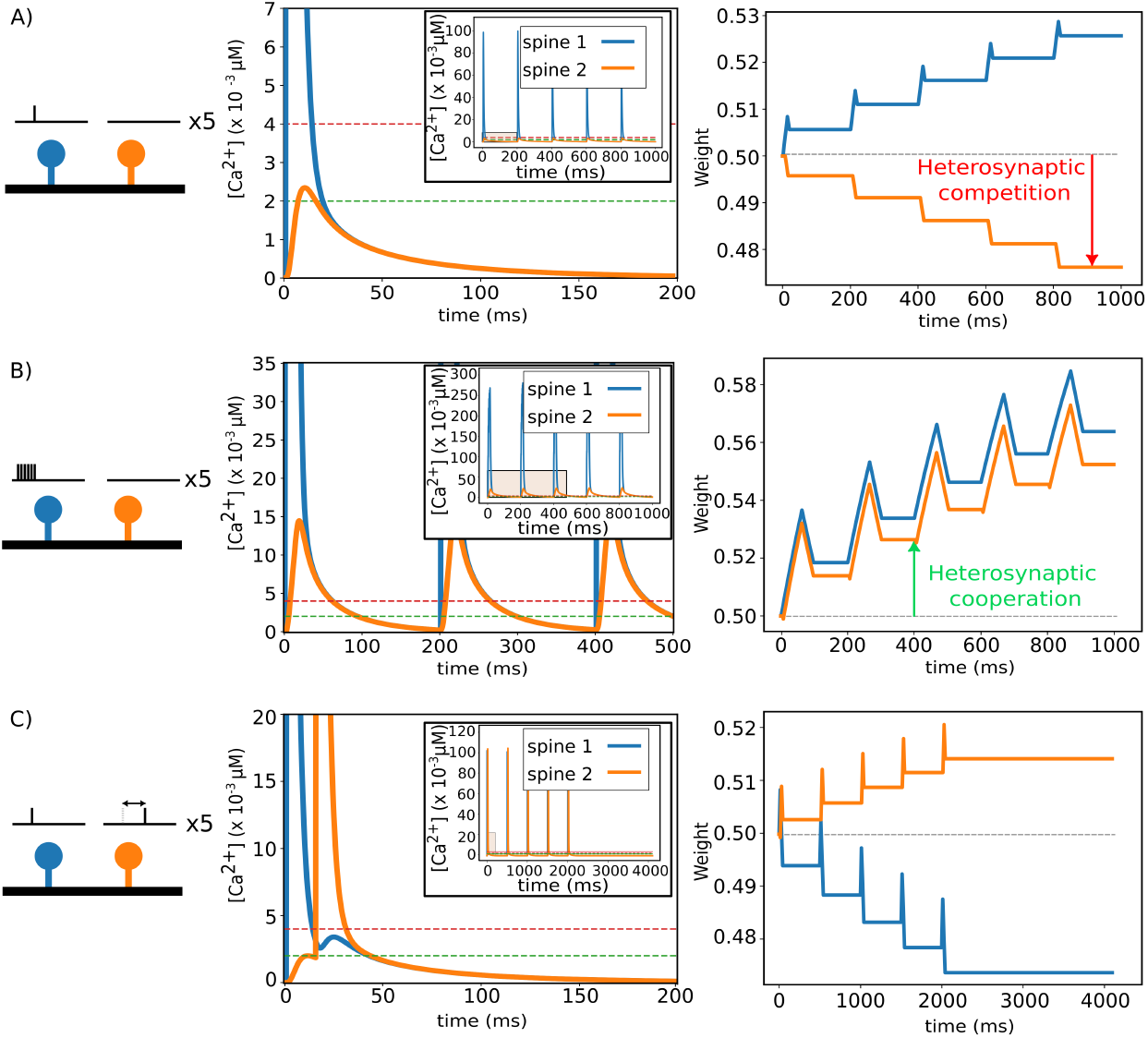
Heterosynaptic effect of different stimulation protocols. Each stimulation protocol is given 5 times. Left: stimulation protocol; Middle: temporal development of calcium concentrations in spines; Right: temporal development of synaptic weights. Blue: Spine 1; Orange: Spine 2. A) Single input stimulation at one spine, while the second spine receives no input signal. B) Burst of inputs at one spine with second spine being unstimulated. C) Single input stimulation of both spines with time difference of 15*ms*. Parameters: θ_*d*_ = 2.10^−3^µM (green dash line), θ_*p*_ = 4.10^−3^µM (red dash line).

**Figure 3:**
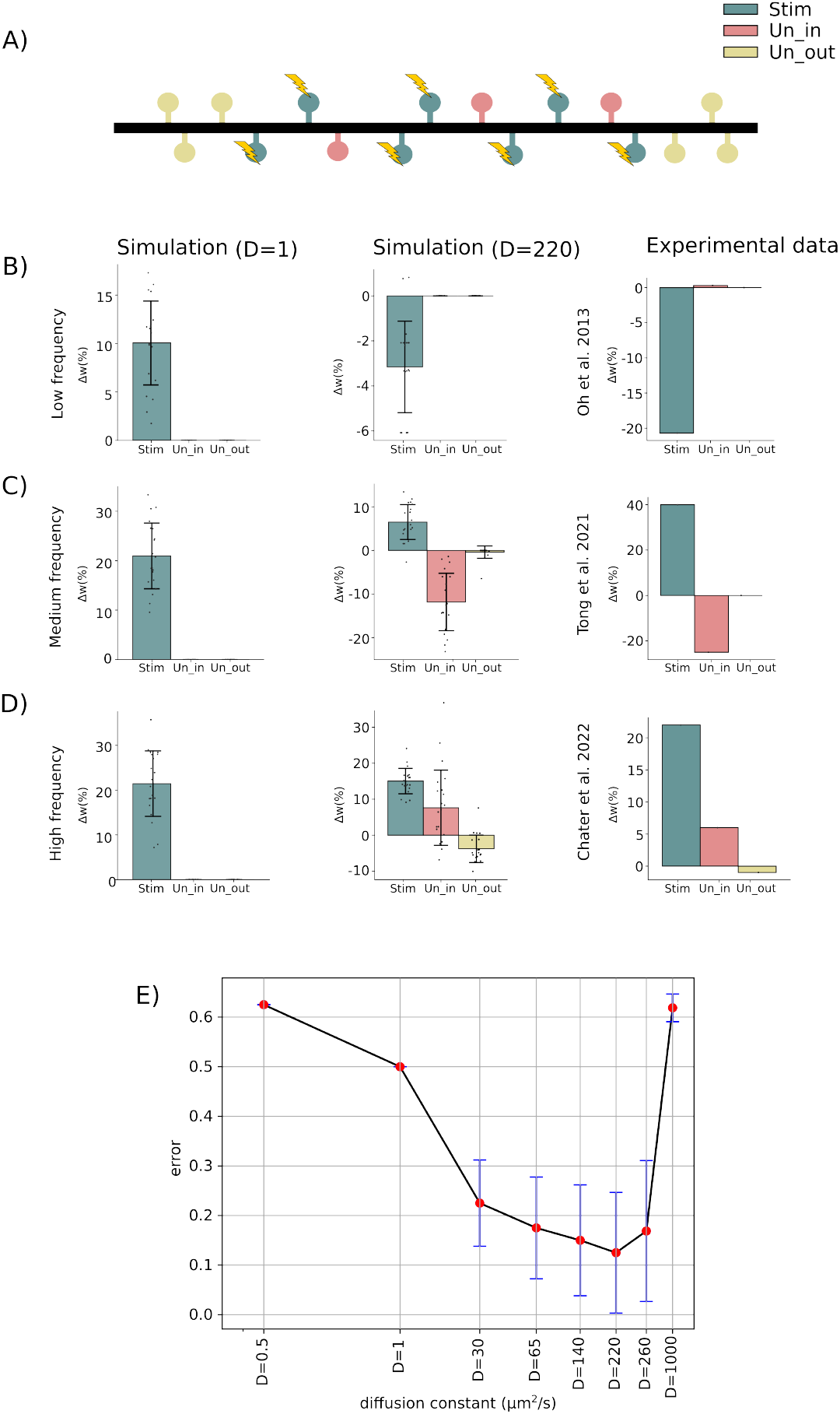
Simulation and experimental results at different frequencies (2, 100, and 150 Hz). **A**) Arrangement of stimulated (in gray) nearby unstimulated (in light red) and distant unstimulated spines (in yellow). **B)-D**) Simulation results with low 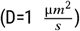 and physiological diffusion constant 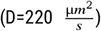 as well as experimental results. **B**) Simulation results with low input frequency, **C**) medium frequency, and **D**) high frequency. **E**) Errors between simulation and experiments shown in B)-D) for different diffusion constants (To see patterns of synaptic weights of these values see Fig. S8). Note that diffusion constants are plotted in logarithmic scale.

Next, we extended our model considering a dendrite with multiple spines to match our model results to various experimental data sets (Chater et al. 2022; Oh et al. 2015; Tong et al. 2021). Note that the experimentally reported plasticity patterns after stimulation show a high degree of variety, lacking a coherent explanation. To integrate all three data sets, we modeled a 80µm long dendrite hosting 16 randomly distributed spines that are governed by calcium-dependent synaptic plasticity (Fig. 3A). All protocols have been repeated for 20 different, random spine configurations, in which initial synaptic weights are randomly chosen representing different initial sizes and volumes of the spines in experiment. A random subset of 7 spines receives stimulation and forms in the following the group of stimulated spines (Stim, gray). We ensure that at least three non-stimulated spines are situated outside the group of stimulated spines on each side (Un_out_, yellow). All further spines are also non-stimulated and form the group of inner neighboring spines (Un_in_, red).

In Oh et al. 2013, it has been shown that a low-frequency uncaging induces synapse-specific spine shrinkage and, thus, likely LTD, while the unstimulated synapses remain unaffected (Fig. 3B). To match this pattern of plasticity, we applied a low-frequency stimulation to the 7 out of 16 randomly distributed spines and also find on average a depression of synaptic weights of the Stim group. Consistent with experimental findings, unstimulated spines maintain their initial state, showing no significant plasticity.

If we increase the stimulation frequency of the targeted stimulated spines, we can qualitatively match the pattern of plasticity from Tong et al. 2021 (Fig. 3C). Here, the average synaptic weights of stimulated spines are potentiated, while we observe LTD at the inner neighboring, unstimulated spines and no significant change at the outer, unstimulated spines.

Further increasing the stimulation frequency at targeted spines enables us to match the experimental results from Chater et al. 2022 (Fig. 3D). Using glutamate uncaging, the authors observed that, after stimulation, the volumes (and accordingly the synaptic weights) of stimulated and inner neighboring, unstimulated spines grow, while the volume of the Un_out_ group decreases.

Note that for all three protocols we show the pattern of plasticity averaged across 20 different spine distributions. For individual distributions, results could vary considerably in the model as well as in experiments (see Fig. S7 and also Chater et al. 2022). Furthermore, the induction of heterosynaptic plasticity is essential for all protocols. If we reduce the effective diffusion constant such that heterosynaptic plasticity will have a negligible effect on the plasticity pattern, model results always show only homosynaptic potentiation and no plasticity at unstimulated spines (left, Fig. 3B-D).

We repeated all protocols with different values of the diffusion constant (D = 1, 30, 65, 140, 220, 260, 300, 350, 400, 440 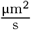) and 20 random configurations each (Fig. 3E). Then, we counted the number of instances where the resulting plasticity patterns matched the plasticity pattern of the corresponding experiment for each spine group (i.e., Stim, Un_in_, and Un_out_, as shown in the last column of Fig. 3B, C, and D), and derived a matching error between 0 and 1. It is evident from Fig. 3E that the lowest error value corresponds to a range of diffusion constants near the calcium diffusion constant. This result supports the model assumption of calcium being one of the main candidates to communicate heterosynaptic plasticity.

After matching the model to experimental data, next, we analyzed the influence of input timing on the plasticity pattern of a system consisting of two spines. Similar to homosynaptic spike-timing-dependent plasticity, we evaluate the temporal window of plasticity, by considering the temporal difference between a spike at spine 1 and a spike at spine 2 and the resulting synaptic weight changes. Note that here we obtain two curves, as we always have two synapses being involved. Fig. 4A, shows the categorized patterns of synaptic weights at different input time-differences. It is evident that increasing γ_d_ enhances the level of depression, thereby shifting the temporal window towards depression for all time differences (green region in Fig. 4A). Similarly, increasing the potentiation coefficient, γ_p_, shifts the temporal window toward potentiation for all time differences (red region in Fig. 4A). There are some intermediate values of γ_d_ and γ_p_ at which the temporal windows are becoming more complex.

**Figure 4.**
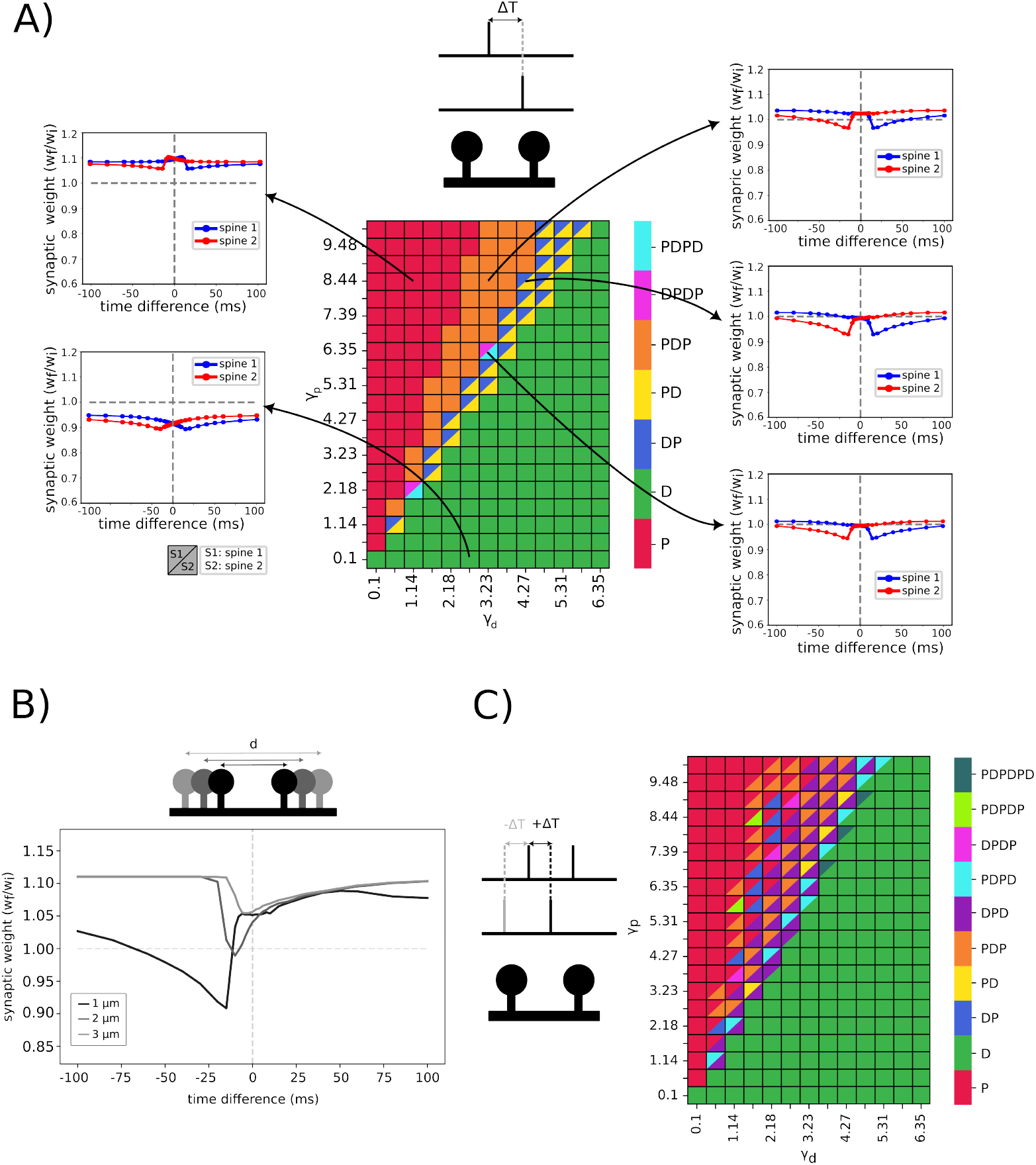
Various curves of heterosynaptic, input-timing plasticity. **A**) Emergent temporal windows after stimulation of two spines at various time differences. Both spines receive one input spike as stimulation. The coefficients γ_*p*_ and γ_*d*_ correspond to the strength of potentiation and depression, respectively. Distance between spines is 1 µm. Letters P and D denote potentiation and depression, respectively. Combinations of these letters indicate the order of occurrence within the total time window from −100 to 100 ms (For calcium-dependent threshold, see Fig. S1, S3). Note that spine 1 is on the left and spine 2 is on the right side of the dendritic branch, and the timing is referenced to the input time of spine 1. **B**) Patterns of calcium-based input timing synaptic plasticity for different inter-spine distance between spine 1 and spine 2 (1,2 and 3 µm). Due to the symmetry between spines 1 and 2, only the synaptic weight of spine 1 is shown. C) Triplet-dependent heterosynaptic plasticity. Emergent plasticity patterns after stimulation of two spines at various time differences with one spine receiving two input spikes and the second spine receiving one spike with a delay. The inter-input interval is 20 ms (For other calcium-dependent parameters, see Fig. S5, S6).

In Fig. 4B, we varied the distance between two spines (1, 2, and 3 µm) and performed the same protocol time-difference protocol. An increased inter-spine distance correlates with a diminished capacity for one spine to influence the state of its neighbors. Consequently, spines situated further apart exhibit reduced heterosynaptic plasticity, tending towards an isolated state (also see **Fig. S2**).

It has been shown that, the traditional pair-based STDP models fail to account for more complex temporal interactions and to explain experimental observations. Therefore, a triplet protocol was suggested to overcome the limitation of canonical pair-based homosynaptic STDP models (Bush and Jin 2012; Pfister and Gerstner 2006). Following this idea, we employed a triplet stimulation protocol involving two input spikes at one spine and a single input spike at a second spine (Fig. 4C). The time differences are calculated relative to the arrival time of the first input spike at spine 1. As anticipated by comparing Fig. 4A and 4C, introducing a third input spike significantly enriches the complexity and diversity of temporal windows of plasticity.

## 4 Temporal sequence selectivity

On the one hand, as shown in Fig. 2 and Fig. 3, dendritic branches are sensitive to the timing and location of synaptic inputs at the spines. Depending on these input features, calcium-based learning can result in distinct signaling patterns within a branchlet. On the other hand, experimental evidence indicates that dendritic branches enable the soma to discriminate the direction of the sequence of inputs - whether it is from the tip of the branch towards the soma (inward sequence), or vice versa as outward sequence (Branco et al. 2010; Fig. 5A). A passive dendritic cable without synaptic plasticity will always result the soma to show the maximum response to the inward sequence. However, for a neuron to be able to discriminate arbitrary input sequences, it should be able to also learn to maximally respond to outward sequences.

**Figure 5.**
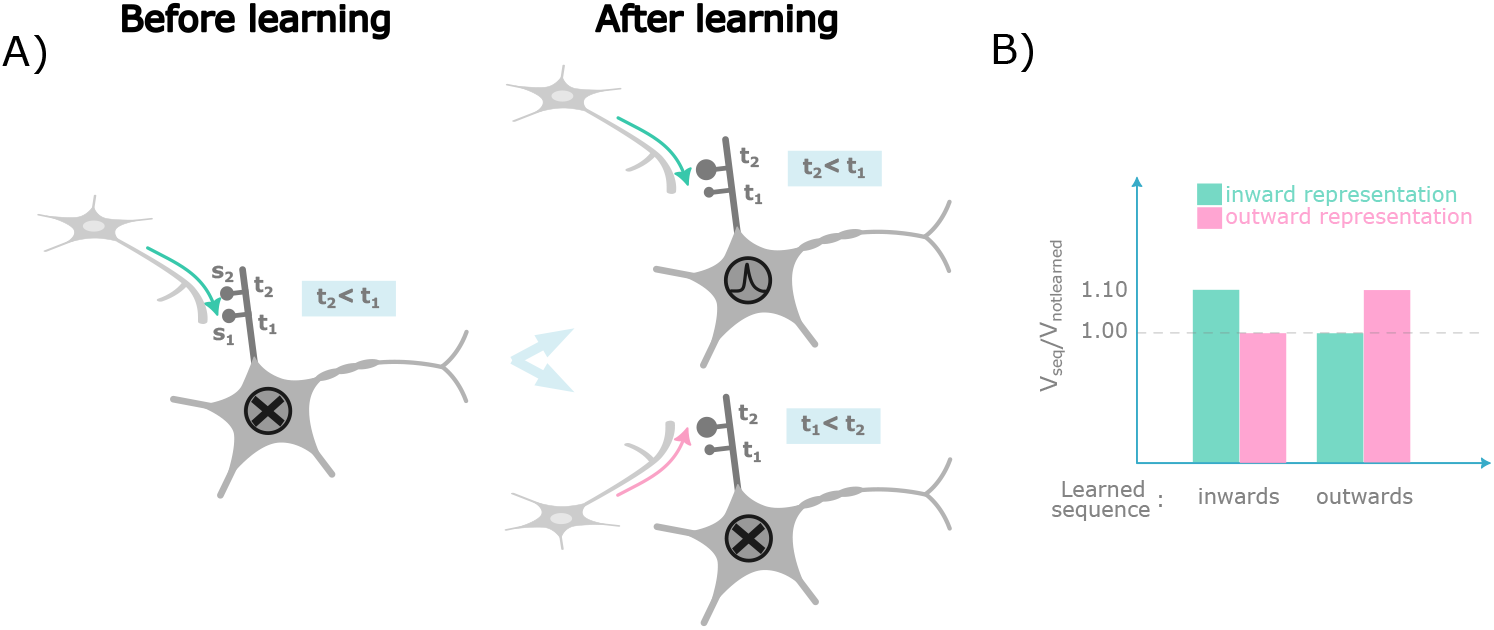
Sequence selectivity via heterosynaptic plasticity. **A**) Illustration of a typical neuron before and after learning an inward sequence. The soma fires in response to the inward sequence but not to the outward sequence. **B**) The ratio of the somatic response presenting the learned versus the non-learned sequence. Left shows the ratios for the protocol in which the inward sequence has been presented during the learning phase, while the right illustrates the ratios when the outward pattern has been presented during the learning phase.

To show that calcium-diffusion-dependent plasticity enables a neuron to learn to discriminate between inward and outward sequences, we introduced a somatic compartment positioned on one end of the dendrite. Hereby, a current flows toward the soma when the membrane potential at the terminal dendritic compartment exceeds the resting potential of the soma. Notably, we did not account for feedback from the soma to the dendrite, neglecting backpropagating effects. The somatic compartment is modeled as a simple integrative unit connected to the terminal dendritic compartment via an axial resistance, R_*s*,axial_. Its electrical properties are defined by its resistance (R_soma_) and capacitance (C_soma_). Here, u_rest_ = − 70mV, τ = R_soma/_C_soma_ = 22ms (Eyal et al. 2016) and R_soma_ R_*s*,axial_ = 5. Note that, (*u*_dend_ *x*_0_), *t* is the voltage at the dendritic compartment being connected to the somatic compartment.

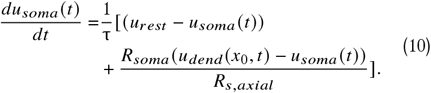

For simplicity, we considered two dendritic spines receiving synaptic inputs through two synapses. These spines were positioned 1µ*m* apart and located proximally to the soma (5µ*m* away).

We first presented the inward input sequence five times to the dendrite (i.e., t_2_ < t_1_) during the learning phase (Fig. 5A). After the learning phase, we presented the inward and outward sequences once, separately, and recorded the somatic membrane potential. We observed that the soma reached its peak potential when the inward pattern was presented (Fig. 5B). Different to a passive, non-plastic dendrite presenting the outward pattern during the learning phase, resulted in the soma exhibiting its maximum response when the outward pattern was presented during the test phase. These findings demonstrate that, by incorporating diffusion-dependent calcium-based synaptic plasticity, our model of a simple dendritic branch can learn different sequences of input signals.

## 5 Discussion and conclusion

We developed a computational model of molecular diffusion dynamics in a piece of dendrite and connected spines. With a diffusion constant that is similar to the one of calcium, our model can match results from several experimental studies of homo- and heterosynaptic plasticity. Our model also shows that given the sensitivity of calcium influx on the timing of presynaptic input spikes, the triggered homo- and heterosynaptic plasticity can lead to complex input-spike-timing-dependent plasticity patterns. This provides a rich repertoire of calcium-based learning rules without the need of postsynaptic spiking.

While our objective was to develop a computational model that balances biological realism with computational practicability, it is essential to recognize the possibility for further improvements. Future improvements could incorporate more intricate and detailed calcium sources, such as calcium-dependent channels or organelles that play a role in synaptic and dendritic calcium dynamics. For example, the rise in cytosolic Ca^2+^ concentration is driven by the influx of extracellular Ca^2+^ through open membrane channels within synapses, but could also be modulated by calcium channels along the dendritic cell membrane (Martucci and Cancela 2022). Intracellular Ca^2+^ release from stores such as the endoplasmic reticulum (ER), mitochondria, and acidic Ca^2+^ stores (i.g lysosomes and endosomes) can also vary intracellular Ca^2+^ concentration and signaling (Martucci and Cancela 2022; Padamsey et al. 2019). Ryanodine receptors (RyRs) are also pivotal Ca^2+^ release channels within the endoplasmic reticulum, contributing to both NMDAR-dependent LTD and LTP under low frequency stimulations (Martucci and Cancela 2022; Segal and Korkotian 2014). RyRs are activated by Ca^2+^ influx and can amplify the effect of the incoming influx of Ca^2+^ signals, potentially lowering the threshold for LTP in neighboring synapses (Padamsey et al. 2019; Segal and Korkotian 2014). Calcium signaling can also occur through store-operated Ca^2+^ entry (SOCE), a mechanism independent of neuronal activity, and facilitate Ca^2+^ influx from the extracellular space. Both RyR-mediated Ca^2+^ release and SOCE play significant roles in shaping synaptic plasticity (Padamsey et al. 2019; Segal and Korkotian 2014). Furthermore, pre- and postsynaptic Ca^2+^ stores are subject to dynamic regulation, influencing the expression of synaptic plasticity (Padamsey et al. 2019). Given more detailed knowledge about the positioning, functioning, and dynamics of such Ca^2+^ stores, our model can be extended to obtain a more complete picture about the functional implication of calcium signals.

The dendritic and synaptic morphology has also a large impact on the observed plasticity patterns. Similar to a study showing the influence of spine density on the diffusion of Cl^−^ ions (Mohapatra et al. 2016), spines can act as barriers for diffusion, reducing the effective diffusion constant, which directly influences the pattern of homo- and heterosynaptic plasticity. Furthermore, changes of the thickness of the spine neck after LTP induction (Kruijssen and Wierenga 2019) can modulate the flux of Ca^2+^ between spine and dendrite, influencing how strongly a synapse can be influenced by heterosynaptic plasticity. Similarly, the positioning of receptors or of the spine apparatus can modulate the flux of Ca^2+^ between spine and dendritic segment (Breit et al. 2018; Noguchi et al. 2005; Rosado et al. 2022). It is worth noting that aging can change the spine morphology and, thus, synaptic plasticity (Bloss et al. 2011). However, in our study, we focused on early developmental stages, during which calcium is less confined to the spine head (Kruijssen and Wierenga 2019; Lee et al. 2016).

Taken together, our results indicate that the intricate, non-linear nature of the spatial and temporal integration of molecular signals introduces a level of complexity that can result in complex patterns of synaptic plasticity. Especially the sensitivity to input spike-timings equips neuronal systems with a large and rich repertoire of learning rules, whose computational potential has yet to be discovered. In a series of studies, one specific realization of such a heterosynaptic input-spike-timing-dependent plasticity rule (Porr and Wörgötter 2006) provides a glimpse into the computational benefits and potential technological applications of heterosynaptic learning mechanisms, like active noise reduction (Möller et al. 2022), control of walking robots (Manoonpong et al. 2007), or learning with optical fibers (Ortín et al. 2023).

## 6 acknowledgments and funding

This work was supported by the German Research Foundation (Deutsche Forschungsgemeinschaft, DFG) through grants SFB1286/C01&Z01, TE 1172/7-1, as well as by the European Commission H2020 grant no. 899265 (ADOPD) and 945539 (HBP SGA3). We thank Arash Golmohammadi Naghshe for his valuable feedback and insightful comments.

## Notes

### Competing Interest Statement

The authors have declared no competing interest.

### Summary of Updates

Section 4 has been added. Figure 5 added. Supplemental files updated.

