## Supplementary information for "Calcium-based input timing learning"

### Supplementary information : Calcium-based input timing learning

Shirin Shafiee [α,β], Sebastian Schmitt[α,β], and Christian Tetzlaff[α,β]
[α] III. Institute of Physics – Biophysics, Faculty of Physics, University of Göttingen, Göttingen, Germany
[β] Group of Computational Synaptic Physiology, Department of Neuro- and Sensory Physiology, University Medical Center Göttingen, Germany

This study investigated the plasticity of synaptic connections under various Calcium-based input timing learning conditions, manipulating different parameters. We measured the inter-spine distances ($\mu$m) and evaluated calcium-related coefficients and thresholds. Our findings revealed distinct plasticity patterns. As discussed in the main text, a triplet pattern of inputs can lead to asymmetrical synaptic weight distributions and more detailed results are represented here.

#### 1 Single input stimulation

Calcium-based input timing patterns for a pair of single-input at distance 1 and 2 $\mu$m are shown in Fig. 1, Fig. 2a and Fig. 2b.

##### 1.1 Distance dependency

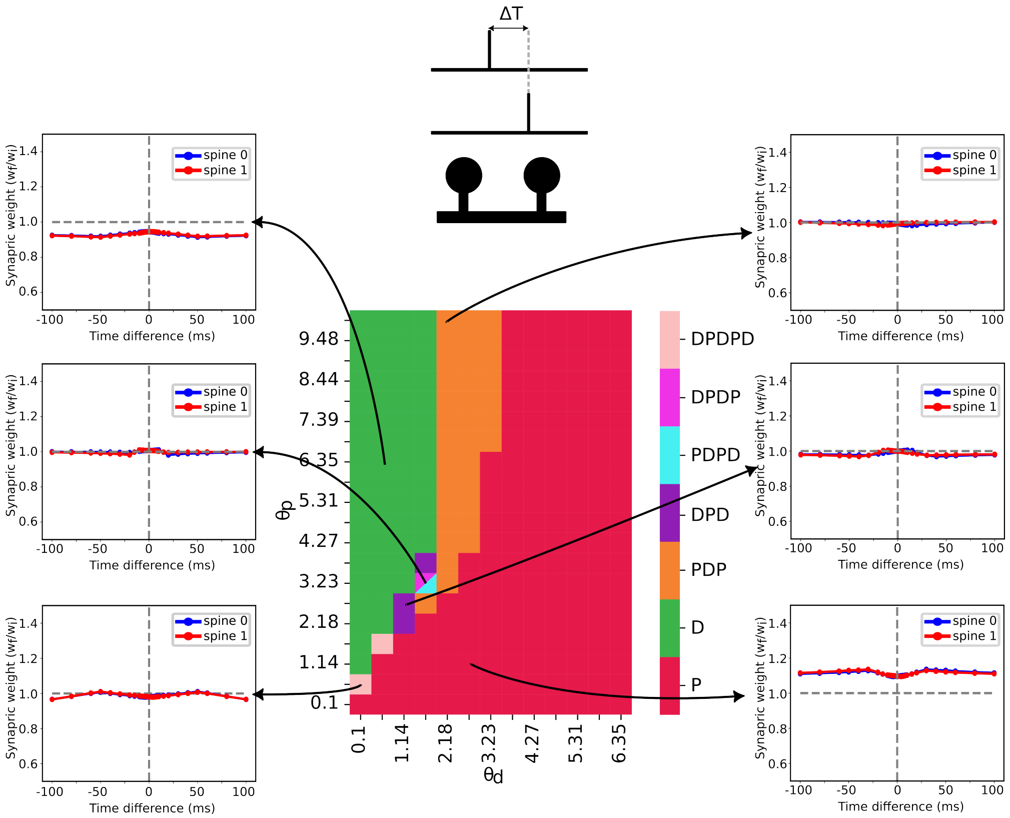


Figure 1: **Two spines at a distance of 1$\mu$m from each other.** Calcium-based input timing patterns are plotted at different potentiation and depression thresholds.

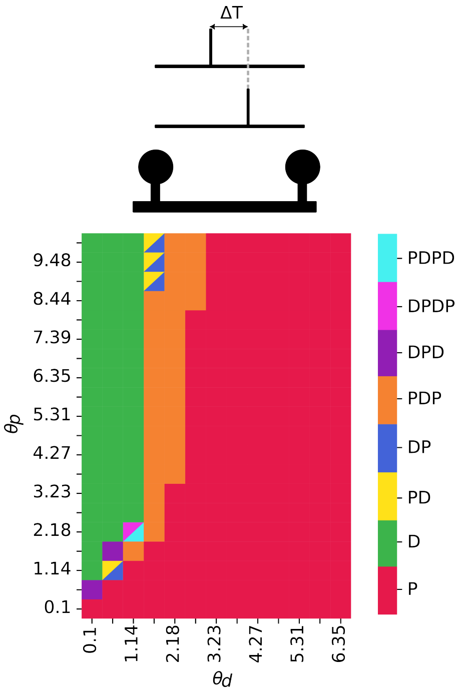


(a) **Calcium thresholds**

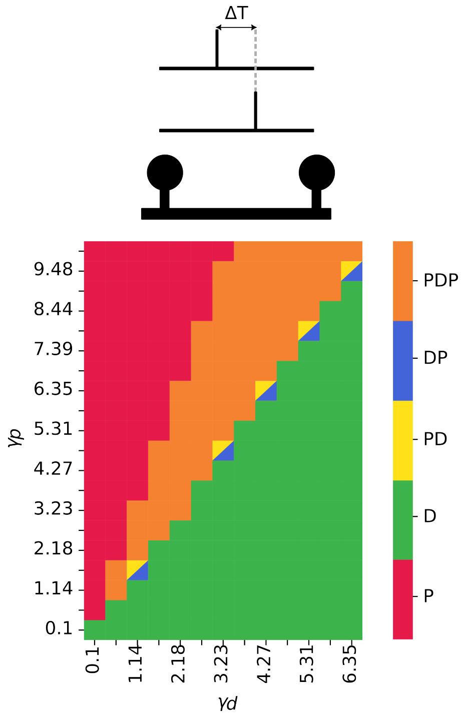


(b) **calcium-related coefficients**

Figure 2: **Two spines at a distance of 2$\mu$m from each other.** Calcium-based input timing patterns are plotted at 2a different potentiation and depression thresholds and 2b calcium-related coefficients.

#### 1.2 Input repetition

Similar stimulation protocols as employed in Section 1 were used, with the exception that each pattern was repeated five times. The results demonstrate that repeated stimulation led to more pronounced changes in synaptic weights, as evidenced by a comparison of Fig. 3 and Fig. 1.

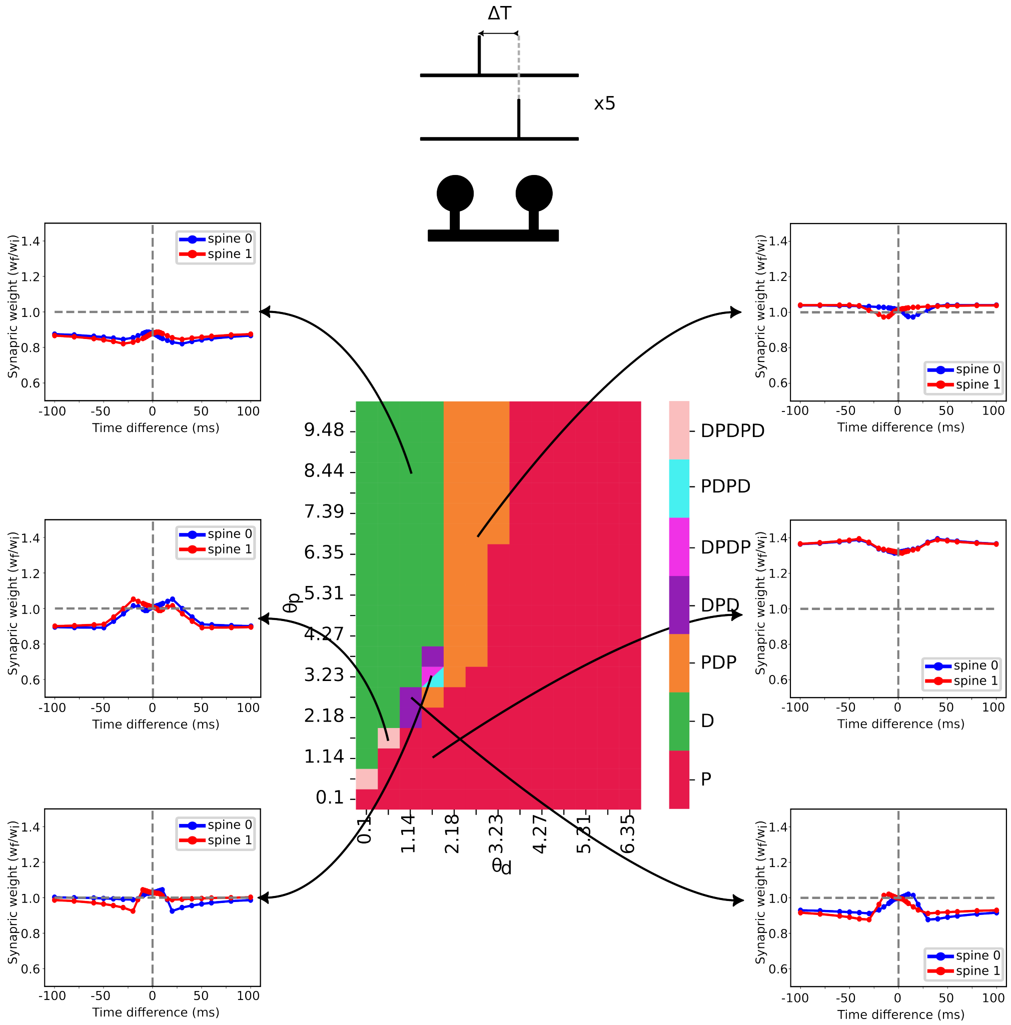


Figure 3: **Calcium-based plasticity patterns reveal enhanced synaptic weight changes with repeated input stimulation**. Calcium-based input timing patterns plotted at a distance of $1\mu$m with varying calcium-dependent thresholds. Stimulation is repeated 5 times with a 200 ms interval between events.

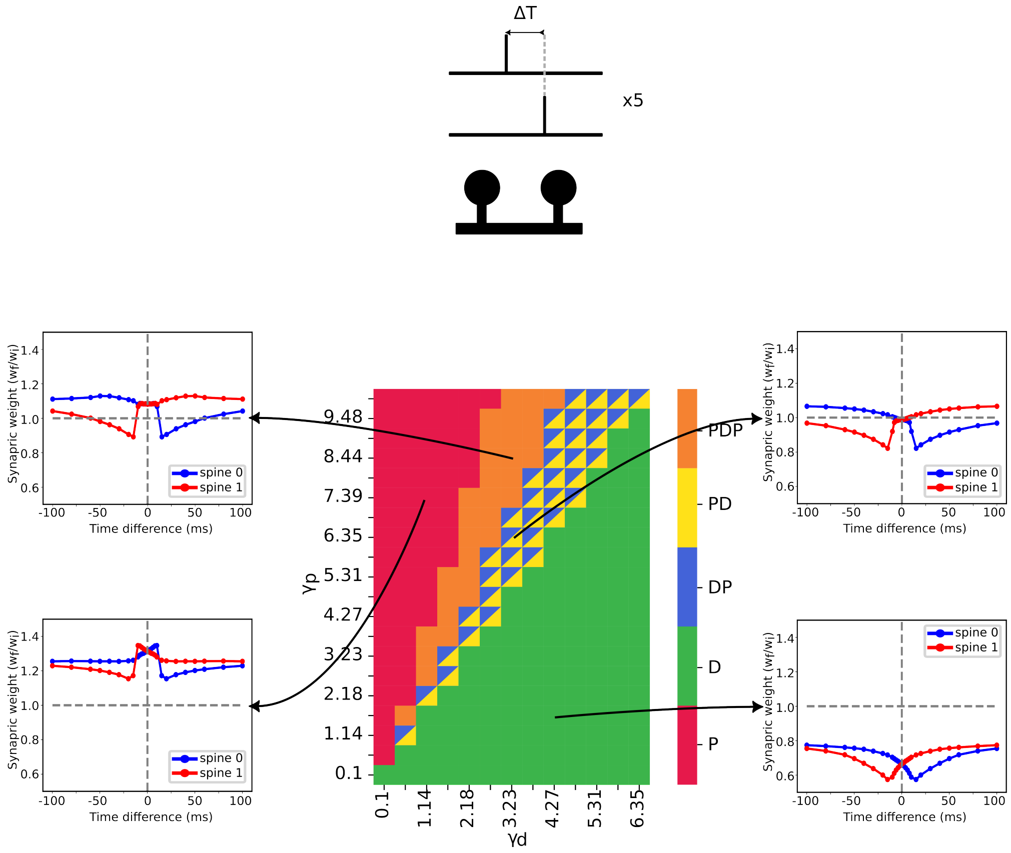


Figure 4: **Calcium-based plasticity patterns reveal enhanced synaptic weight changes with repeated input stimulation**. Calcium-based input timing patterns plotted at a distance of 1$\mu$m with varying calcium-dependent coefficient. Stimulation is repeated 5 times with a 200 ms interval between events.

### 2 Triplet inputs

As demonstrated in the main study, the triplet input protocol alters the temporal dynamics of calcium signal integration, leading to subsequent changes in synaptic weights and, consequently, the Calcium-based input timing patterns (Fig. 5 and Fig. 6).

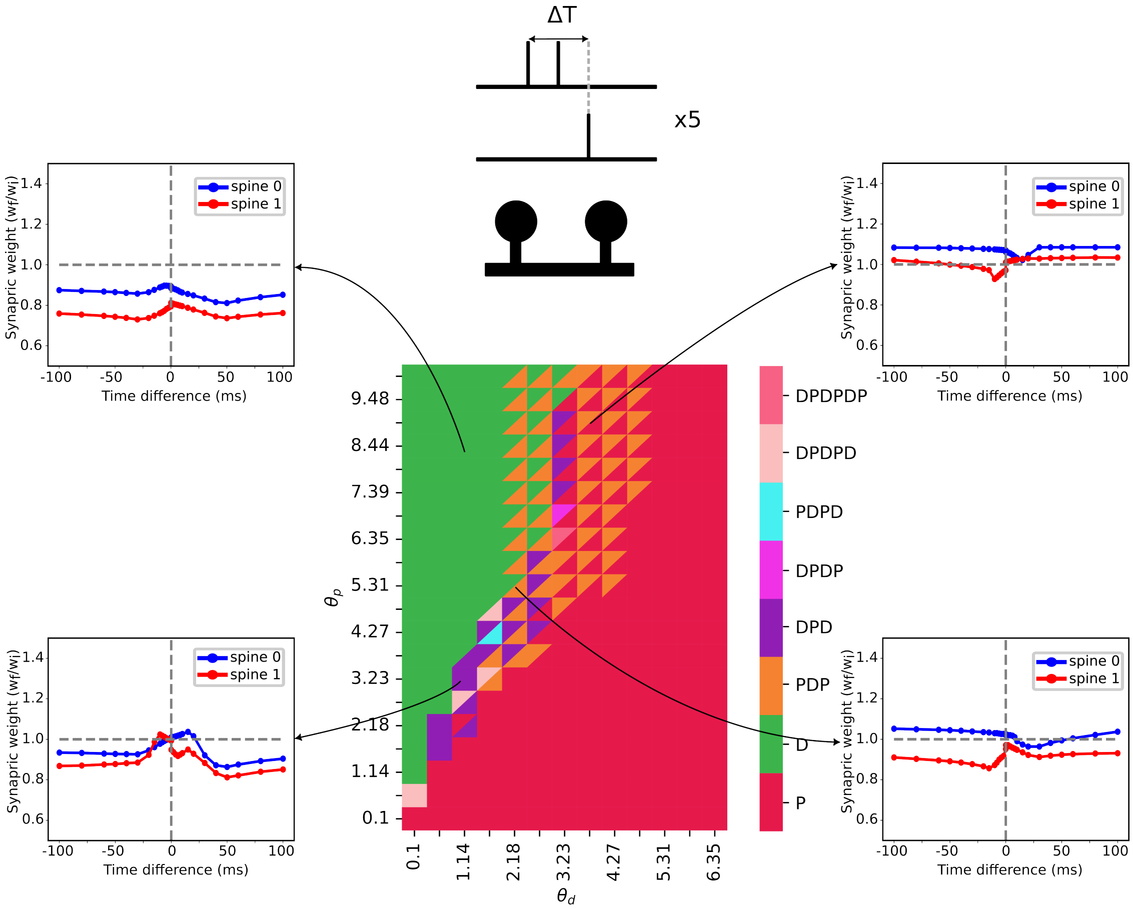


Figure 5: **Triplet-dependent heterosynaptic plasticity at different calcium-dependent threshold.** Emergent plasticity patterns after stimulation of two spines at various time differences with one spine receiving two input spikes and the second spine receiving one spike with a delay. The inter-input interval is 20 ms and inter-spine distance is $1\mu$m.

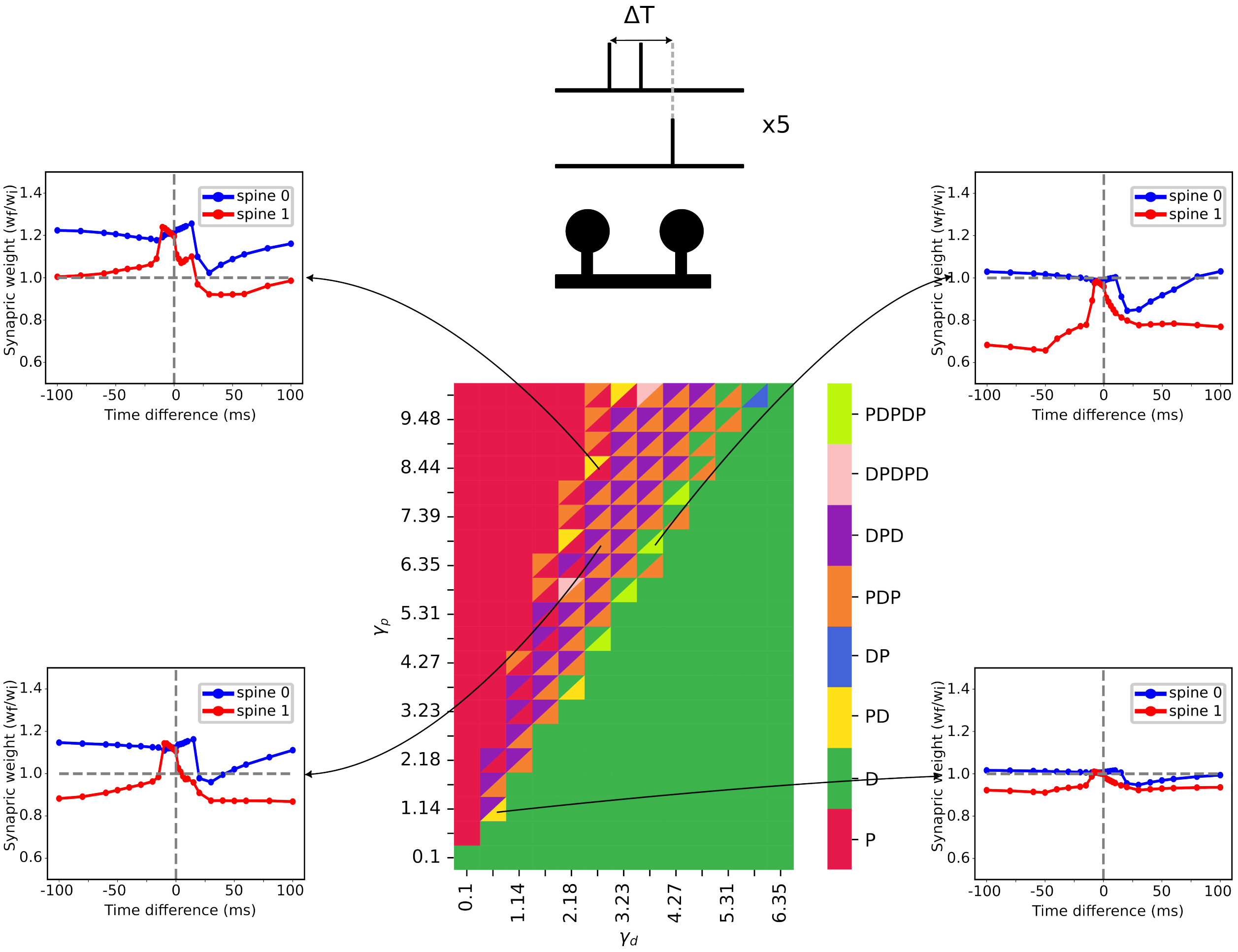


Figure 6: **Triplet-dependent heterosynaptic plasticity at different calcium-dependent coefficient.** Emergent plasticity patterns after stimulation of two spines at various time differences with one spine receiving two input spikes and the second spine receiving one spike with a delay. The inter-input interval is 20 ms and inter-spine distance is $1\mu$m.

### 3 Multi-spine stimulation

As previously discussed in the main study, the emergent patterns of synaptic weight modification are highly sensitive to the specific characteristics of the system, particularly the relative positions of spines on the dendritic branch. Here, three distinct spine configurations under varying stimulation frequencies investigated.

For instance, in the first configuration depicted in Fig. 7, three stimulated spines are positioned in close proximity. This clustered arrangement may contribute to the formation of stronger and higher synaptic weights at higher frequencies compared to stimulated spines in other configurations.

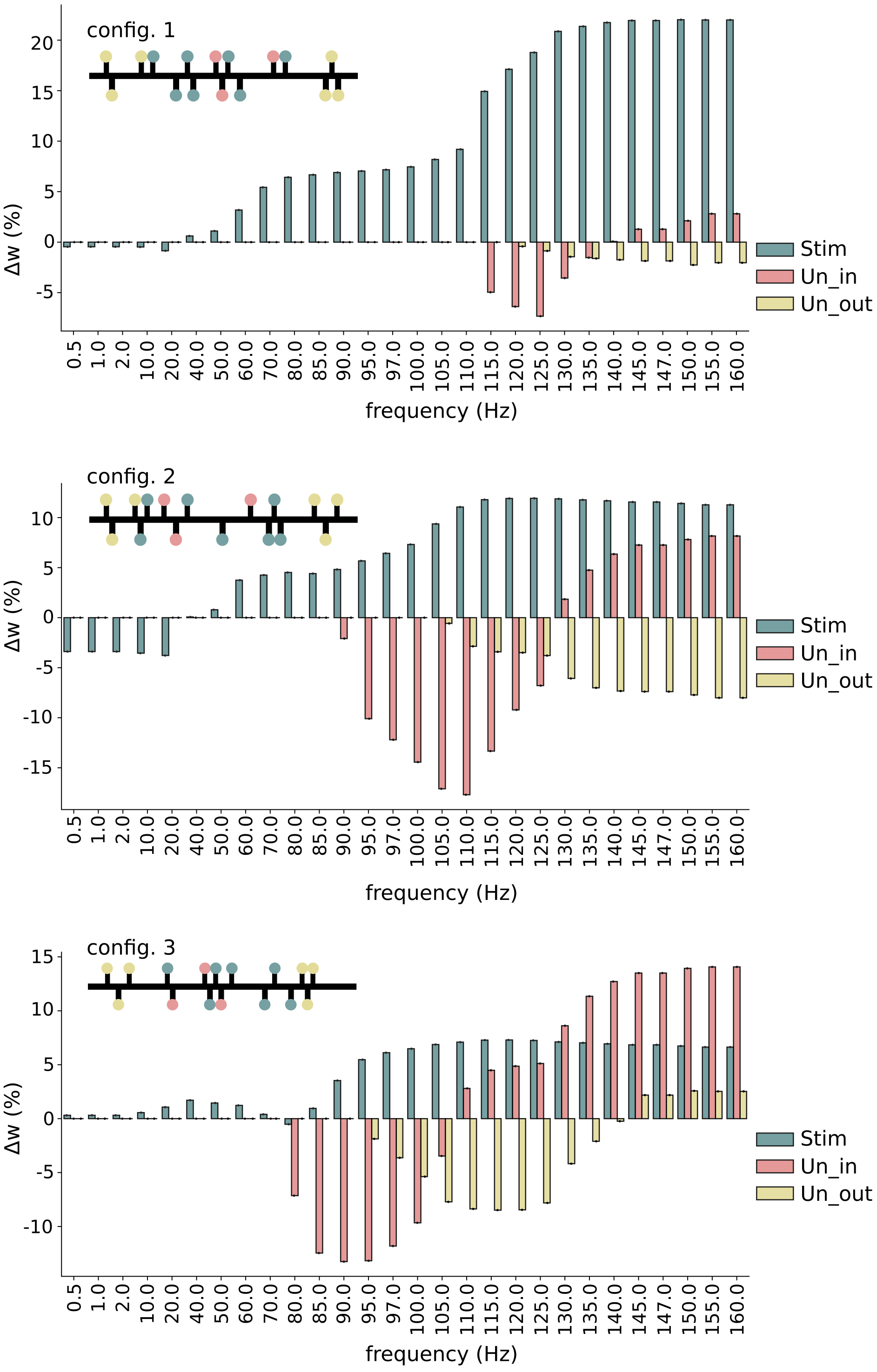


Figure 7: **Multi-spine stimulation**. Three distinct spine configurations with varying initial synaptic weights were considered. The average change in synaptic weights is plotted against different stimulation frequencies.

#### 3.1 Different diffusion constants

The same simulation as presented in Fig. 3 of the main study was replicated using four different diffusion constants: 1, 65, 220, 300, and 400 $\frac{\mu m^2}{s}$. The following results were obtained.

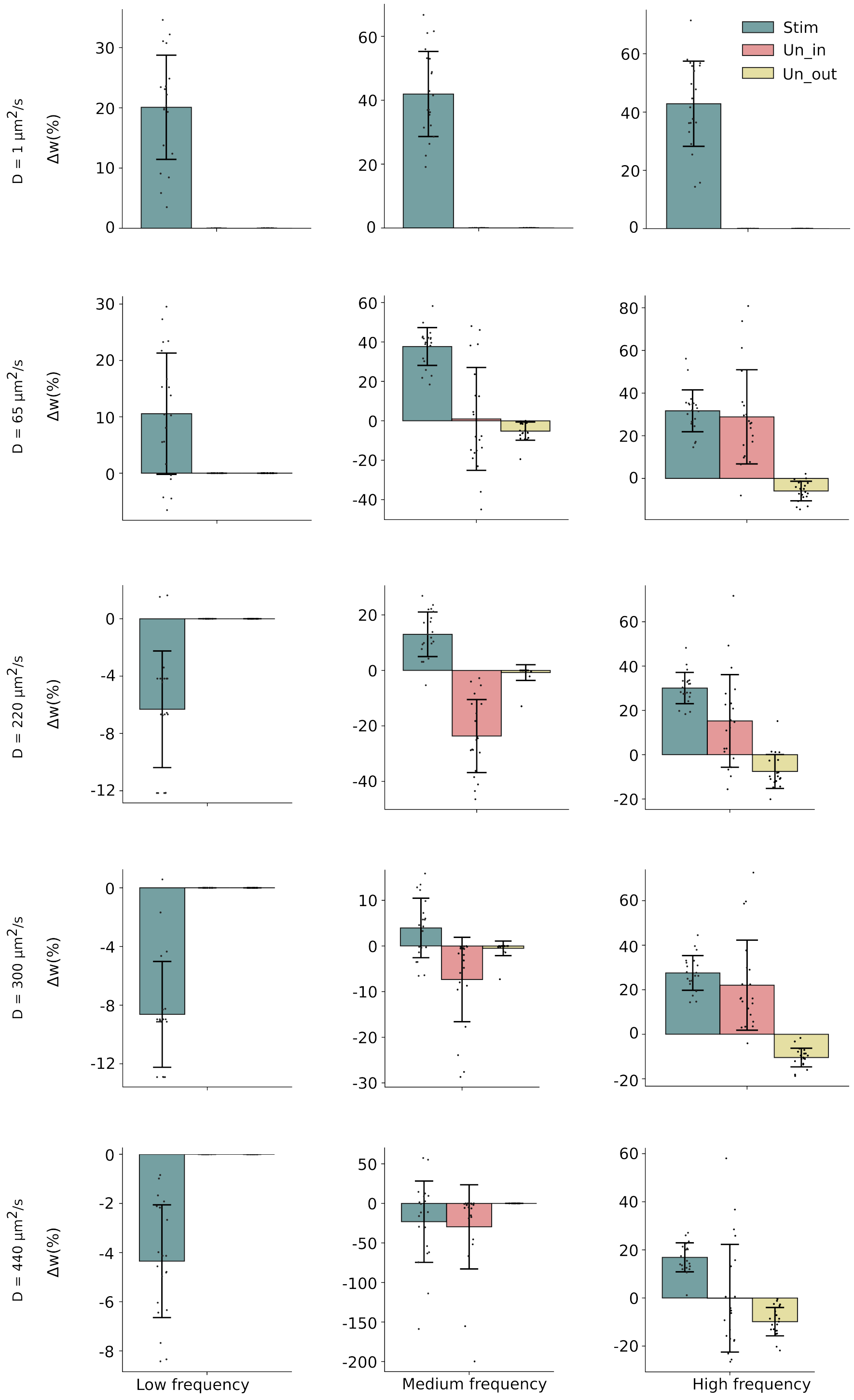


Figure 8: **Simulation results at in different diffusion constants and different frequencies (2, 100, and 150 Hz)**. Simulation results with low (D=1) and physiological diffusion constant (D=220) . Simulation results with low input frequency (first column), medium frequency (second column), and high frequency (third column).

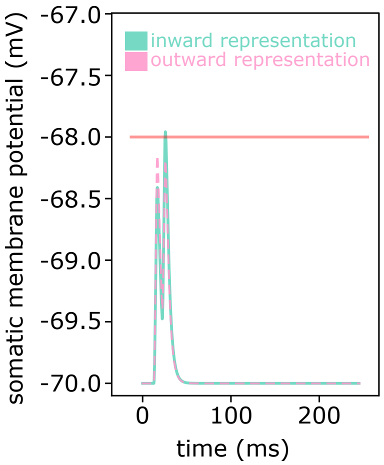


Figure 9: **Sequence selectivity via heterosynaptic plasticity.** Somatic membrane potential after showing inward (green) and outward (pink) sequences. If a spiking threshold of $V_{th} = -68.0$mV is considered, the soma fires an action potential following the presentation of the inward pattern.

### 4 Supplementary methods

#### 4.1 Numerical consideration and steady state

Using forward Euler(FE) scheme forces us to have a limitation choosing every $\Delta x$ and $\Delta t$, as they depends to each other. Applying Von Neumann stability analysis for diffusion equation gives us the limitation for $\Delta x$ and $\Delta t$, e.g $\Delta t \leq \frac{1}{2}\frac{(\Delta x)^2}{D}$ which $D$ is the diffusion coefficient. Avoiding such stability problems, we can use 'Backward Euler' method which is unconditionally stable and robust and does not suffer from numerical instability like the explicit Euler method . In this case, we have the flexibility to select the grid parameters $\Delta x$ and $\Delta t$ independently, without concerns regarding instability arising from their combination. Although the backward Euler method has the same numerical accuracy as the forward Euler method and it does not suffer instability, it has more computational expense [1].
